# Genome characteristic of *Bordetella parapertussis* isolated from Iran

**DOI:** 10.1101/2022.01.05.475152

**Authors:** Azadeh Safarchi, Samaneh Saedi, Chin Yen Tay, Binit Lamichhane, Masoumeh Nakhost Lotfi, Fereshteh Shahcheraghi

## Abstract

Pertussis also known as whooping cough is a respiratory infection in humans particularly in infants and usually caused by *Bordetella pertussis*. However, *Bordetella parapertussis* can also cause a similar clinical syndrome. During 2012 to 2015, from nasal swabs sent from different provinces to the pertussis reference laboratory of Pasture Institute of Iran for pertussis confirmation, seven *B. parapertussis* isolates were identified by bacterial culture, biochemical tests, and the presence of IS*1001* insertion in the genome by real-time PCR. Furthermore, the expression of pertactin (Prn) as one the major virulence factor for bacterial adhesion was investigated using western blot. Moreover, the genomic characteristic of one recently collected isolate, IRBP134, from a seven-month infant was investigated using Illumina NextSeq sequencing protocol. The results revealed the genome with G+C content 65% and genome size 4.7 Mbp. A total of 81 single nucleotide polymorphisms (SNPs) and 13 short insertion and deletions were found in the genome compared to the *B. parapertussis* 12822 as a reference genome showing ongoing evolutionary changes in our isolate. A phylogeny relationship of IRBP134 was also investigated using global *B. parapertussis* available genomes.

## Introduction

*Bordetella parapertussis* is a Gram-negative bacterial pathogen that colonise in the respiratory tract and cause the vaccine-preventable disease whooping cough. However, the severity of the disease caused by *B. parapertussis* is thought to be shorter in duration and milder than *B. pertussis*, the main responsible pathogen for pertussis in human [1, 2]. Comparative genome analysis of *Bordetella* species revealed that *B. pertussis* and *B. parapertussis* were independently evolved from *B. bronchiseptica*-like ancestors [3]. *B. parapertussis* diverged to two distinct lineages; one cause whooping cough in infants and the other infects sheep [1].

*B. parapertussis* and *B. pertussis* share same virulence factors including pertactin (Prn), dermonecrotic toxin, filamentous haemagglutinin (Fha) and adenylate cyclase [4]. However, the pertussis toxin (Ptx) is only expressed in *B. pertus*sis, since the *ptx* operon in *B. parapertussis* is dysfunctional due to the mutation in the *ptx* promoter and coding region [3]. In addition, unlike *B. pertussis, B. parapertussis* isolated from human is oxidase negative and harbour IS*1001* insertion element in the genome [3] which can differentiate the clinical isolates from *B. pertussis* (IS*481*) using rt-PCR in diagnostic laboratories [5]. Therefore, if only the presence of IS*481* is investigated for pertussis confirmation resulting false negative pertussis cases and leading to the low isolation rate of *B. parapertussis*, although most of the diagnosis commercial kits target both IS*481* and IS*1001* simultaneously. There are few reports of widespread infection of *B. parapertussis* [6]. In Iran, we reported increase in the number of B. pertussis isolates in recent years and investigated their genomics [7-10]. However, reports showed the lower isolation rate of *B. parapertussis* compared to *B. pertussis* [11].

Unlike *B. pertussis*, there are a few studies investigating the genomic and proteomic characteristics of *B. parapertussis* or analysing recently collected isolates [3, 12-15]. Here, we investigate the expression of pertactin (Prn) as one of the important immunologic antigens that are responsible in the bacterial adhesion to host cells and are involved in the acellular pertussis vaccine. Furthermore, the genomic characteristic and microevolutionary changes of one recently isolated *B. parapertussis* in Iran was analysed using whole genome sequencing approach and compared with global isolates.

## Material and method

### Bacterial isolation and identification

.During 2012 to 2015 when the pertussis epidemic was reported in Iran [7] nasal swabs of pertussis suspected patients were sent to pertussis reference laboratory of Pasture Institute of Iran for bacterial isolation and infection confirmation. Briefly, samples were cultured on Regan-Lowe medium containing charcoal agar and 10% defibrinated sheep blood and incubated at 37 °C for 72. *B*. para*pertussis* isolates were confirmed by a combination of colony morphology, Gram stain and conventional biochemical tests such as oxidase and real-time PCR. DNA extraction was performed using High Pure PCR Template Preparation Kit (Roche Diagnostics GmbH, Mannheim, Germany) and real-time PCR was performed by targeting IS*481*, IS*1001* and IS*1002* with designed primers [9] to confirm the presence of IS*1001* for *B. parapertussis* as recommended by WHO [5].

### Western Blot

Western blot analysis was performed to investigate the expression of pertactin in our isolates. Briefly isolates were suspended in phosphate-buffered saline (PBS) and boiled in 55 °C for 30 min after bacterial subculture on Bordet Gengou agar with 15% sheep blood at 37 °C for 72 h. *B. pertussis* strain Tohama I (Gene Bank accession number BX470248) and Klebsiella (ATCC 13883) were used as a positive and negative control respectively. Bacterial proteins were separated by 10% sodium dodecyl sulphate polyacrylamide gel electrophoresis (SDS-PAGE) (Bio-Rad). After electrophoresis, the proteins were transferred to polyvinylidene fluoride (PVDF) membranes at 300 V for 1 hr. The membranes were blocked with skim milk in PBS for overnight. The 220-kDa Fha protein and 69-kDa prn was detected using a mouse anti-Prn antibody (NIBSC, UK), then incubated with a horseradish peroxidase (HRP)-conjugated anti-mouse antibody. After a final wash, membranes were developed with Metal Enhanced DAB Substrate (Thermo Fisher Scientific, Waltham, MA, USA).

### Whole genome sequencing

*B. parapertussis* isolate IRBP134 collected in autumn 2015 from the nasal swab of a fully vaccinated 7-month-old female baby was selected for whole genome sequencing. DNA was extracted and purified from pure culture as described previously [16]. DNA libraries were prepared with the insert size of 150 bp paired-end using NexteraXT DNA kit (Illumina) and sequenced on the Nextseq (Illumina) with a minimum coverage of 150-fold. The raw reads were submitted to the GeneBank database under the Biosample number SAMN18214790.

### Bioinformatics

*de novo* assembly and genome annotation were performed as described previously [7, 16]. Assemblies were also submitted to CRISPRfinder and Phaster for CRISPR (clustered interspaced short palindromic repeats), and prophage prediction in the genome respectively [17, 18]. Reads were mapped against *B. parapertussis* strain 12822 (GenBank: NC_002928.3), that is used as a reference genome in most studies, and SNPs and indels were detected as previously described [7, 16]. Virulence-associated gene analysis and the multilocus sequence typing (MLST) were performed using the *Bordetella* Bacterial Isolates Genome Sequence Database (BIGSdb) at https://pubmlst.org/bordetella. The maximum parsimony algorithm was used to construct phylogenetic tree by MEGA7 [19] Tree-Bisection-Reconnection (TBR) was used to search optimal trees. Bootstrap analysis was based on 1000 replicates and *B. parapertussis* 12822 used as reference genome.

## Results and Discussion

### Detection of *B. parapertussis* in clinical samples

During 2012 to 2015 from 4923 swabs sent to Pasture pertussis reference laboratory, seven *B. parapertussis* isolates were confirmed of which four isolates collected from unvaccinated infants with the age of 2-month-old or less and three were collected from fully vaccinated patients older than 6-month-old (Table-1). Tehran as a capital and Eastern Azarbayjan, a north-western province, each had three isolates. These two provinces had the most *B. pertussis* isolates in recent years as reported previously [7]. During 2012-2015 around 112 *B. pertussis* isolates collected from different provinces in Iran [7] showing low isolation rate of *B. parapertussis* in the country with 50 years whole cell pertussis vaccination history. In recent years, there are some reports showing the increase of pertussis cases caused by *B. parapertussis* especially in countries with acellular pertussis (ACV) immunisation program [6, 12, 20, 21]. ACV used since 1990s in most developed countries for immunisation due to the reported side effects of WCV and usually contained three (Ptx, Prn, and Fha) or five components (additional fimbriae Fim2 and Fim3) [22]. The increase in the *B. parapertussis* isolation might be due to the fact that it might have better fitness under the ACV pressure since it does not express Ptx, as the major virulence protein that are involved in all types of one to five component ACVs [23].

**Table 1.** Details of *B. parapertussis* isolates collected in Iran during 2012-2015

| Isolate identification code | collected in | Age | Province | gender | pertussis vaccination status of patients | antibiotic therapy | symptoms |
| --- | --- | --- | --- | --- | --- | --- | --- |
| IRBP8 | Apr. 2012 | 6 years | Eastern Azarbayjan | Female | Pos | Neg | Cough |
| IRBP125 | May-12 | 43 days | Eastern Azarbayjan | Female | Neg | Neg | Cough, vomiting |
| IRBP696 | Sep. 2012 | 44 days | Tehran | Female | Neg | Pos | Cough |
| IRBP1079 | Jan. 2013 | 45 days | Tehran | Female | Neg | Pos | Cough, vomiting |
| IRBP1161 | Feb. 2013 | 2.5 month | Tehran | Female | Neg | Pos | Cough |
| IRBP1396 | Mar-2014 | 7 months | Eastern Azarbayjan | Female | Pos | Pos | Cough |
| IRBP134 | Nov. 2015 | 7 months | Khorasan Razavi | Female | Pos | Neg | Cough, vomiting |

### Prn expression

Pertactin is an important surface antigen in *B. pertussis* and *B. parapertussis* as an adhesion factor that is also included in ACVs. *B. parapertussi* isolates that do not express pertacin were reported in France in recent years [12, 24]. Since 2007, the majority of collected *B. parapertussis* isolates 94.3% in France have not expressed Prn suggested to be due the ACV vaccine pressure [25]. This phenotype is caused by the deletion of one Adenine in region I of the *prn* gene (position 988, 12.1%, 4/33) or a Guanine in region II (position 1895, 75.8%, 25/33) both of which lead to a stop codon [25]. It is shown that prn-negative *B. pertussis* have better fitness under ACV pressure [26]. The emergence of Prn negative *B. parapertussis* isolates could be a global concern especially in countries with ACV immunisation program since this phenotype can easily escape vaccine pressure by not expressing Ptx and Prn as two main components of ACVs. We previously showed that no mutation or disruption was found in the *prn* gene in current circulating predominant *B. pertussis* isolates in Iran and no prn negative isolate was reported from Iran with WCV immunisation program [7]. Here, western blot was carried out to investigate the expression of Prn in our *B. parapertussis* isolates that showed all seven isolates express Prn and confirmed the WCV immunisation did not affect phenotype evolution of this species in Iran.

### General genome features

We sequenced one recently collected *B. parapertussis* isolate, IRBP134, from fully vaccinated infant. The Nextseq sequencing generated 6897792 paired reads with GC content 65% and coverage rate 231. De novo assembly generated 72 scaffolds with genome size 4720964 bps with N50, 106547 (Figure-1).

**Figure-1:**
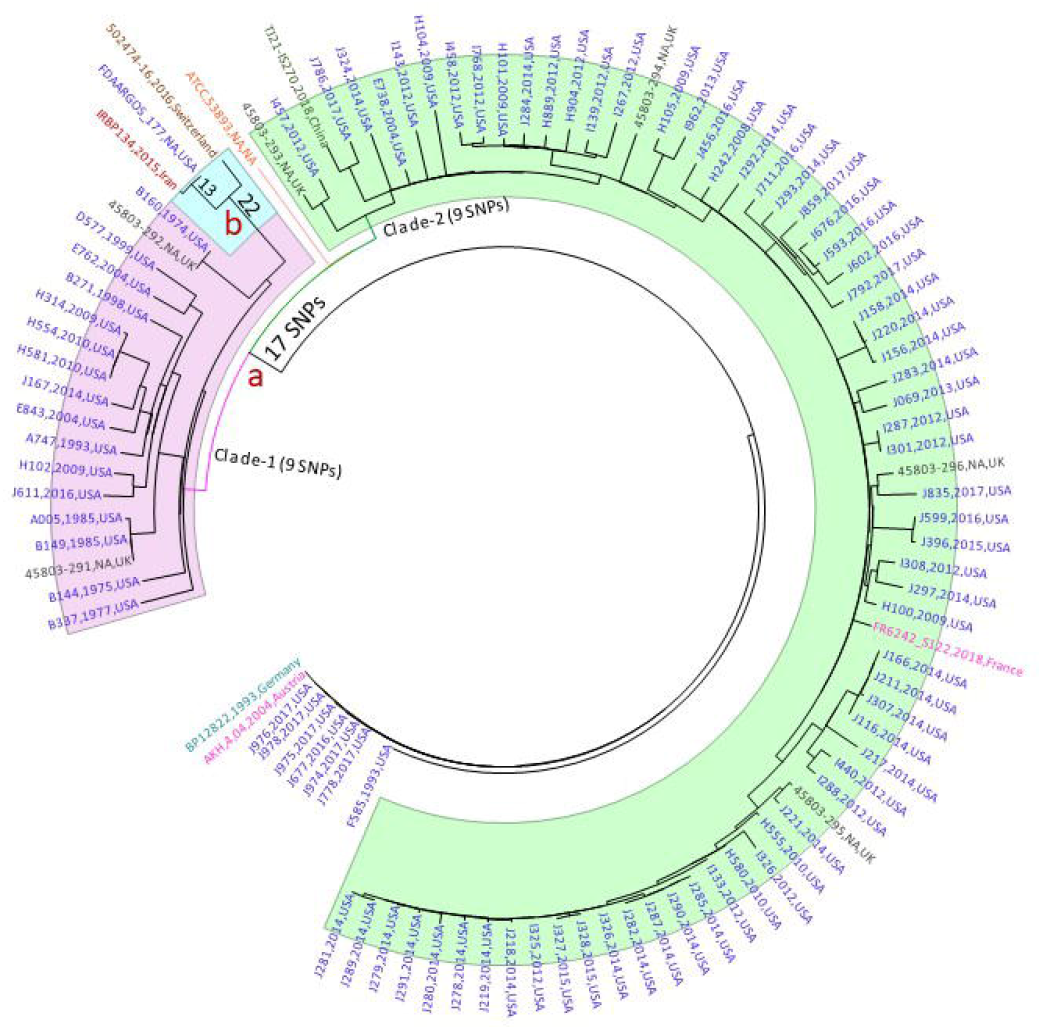
Phylogenetic relationships of Iranian *B. parapertussis* isolate (IRBP134) with global isolates. Maximum parsimony phylogeny using 895 SNPs from a total of 104 *B. parapertussis* isolates from different countries, mainly from the USA: Blue, and other countries such as the UK, France, China, Austria, and Switzerland (details of the isolates are available in Supplementary File-3). *B. parapertussis* strain 12822 was used as a refence. **a)** Majority of isolates (95) carried alle 6 of *brkB* as a virulence associated gene and clustered in two clades as clade 1 and clade 2 each with nine unique SNPs. **b)** The Iranian isolate IRBP134 located in clade 1 which mostly consists of isolates collected before 2000 and make a subclade with two isolates from Switzerland and the USA with 22 SNPs. IRBP134 separated from the USA isolate, FDAARGOS177, with two non-synonymous SNPs located in BPP2476 and BPP3004. The majority of more recently isolates are in Clade −2.

The genome annotation showed the total of 4620 potential coding sequences and 55 RNA including 63 tRNA and one large subunit ribosomal RNA and one small subunit of ribosomal RNA.

PHASTER tool was used to identify phage region in the genome and showed one incomplete phage regions with average size 9.7 kb and GC content 68.17% encoding 11 proteins. The sequence of potential prophages in the genome is identified and categorized as intact, incomplete, or questionable based on the identity score [18].

CRISPR (Clustered Regulatory Interspersed Short Palindromic Repeats) systems were first discovered in *E. coli* in 1987 and later in other species. It is based on the generating specific CRISPR RNA (crRNA) which target invasive RNA/DNA sequences and cleave it into multiple smaller sequences by the endonuclease activity of CRISPR-associated (cas) proteins [27]. Based on the CRISPR/cas database [28], seven CRISPR sequence were found in the genome which is consistent with the average number of CRISPR sequences in other submitted *B. pertussis* isolates in the databases (Supplementary File-1). Furthermore, two genes for CRISPR-associated (cas) proteins (Cas3_1_I and Cas3_0_I), belong to class I Cas proteins, were identified in the genome (Supplementary File-1). We aligned the nucleotide sequences of these two identified Cas proteins in NCBI database using BLASTn tool and found the sequences are identical with *m*fd gene in *B. parapertussis* encoding transcription-repair coupling factor (Mfd) that associates elongation transcription complexes in bacteria and helps RNA polymerase to finish the transcription [29]. Therefore, the identified Cas-associated protein in our genome need to be investigated further to be confirmed as Cas protein since in the CRISPRcas database there was no Cas-related protein to be identified for this specie.

### Microevolutionary analysis

Studies showed pertussis vaccines using *B. pertussis* as vaccine seeds can protect body against *B. parapertussis* as well [5]. There are numerous reports showing allelic variation, genome reduction and ongoing microevolutionary adaptation in currently circulating *B. pertussis* around the word especially in countries switched to acellular vaccine [16, 30-32]. Vaccine pressure particularly the switch from whole cell vaccine to acellular vaccine was one of the main reasons for pathogen adaptation [33-35]. Since *B. parapertussis* cause pertussis with milder symptoms, they isolated and reported less than *B. pertussis* and there are very few studies investigating the adaptation of clinical *B. parapertiussis* isolates [3, 6, 12].

Genotyping of major virulence associated genes revealed IRBP134 had similar virulence associated genes of the reference genome and only exception is in the *brkB* gene that like other *B. parapertussis* isolates [24], it carries allele 6 of *brkB* gene, encoding *Bordetella* serum resistance (Supplementary File-2). This allele variation is a result of a non-synonymous SNP changing polar Threonine to nonpolar Alanine in BrkB in position 742 of an immunologic protein of *B. pertussis*. The virulence associated gene, *brkB*, encodes a cytoplasmic membrane protein called *Bordetella* resistance serum killing. It plays an important role in *B. pertussis* as a virulence factor mediating adhesion of bacterium to the host cell and is also shown to be expressed in *B. parapertussi*s [36]. The multilocus sequence typing (MLST) analysis according to 7-gene scheme [37] shows IRBP134 belonged to ST19 as it is reported for *B. parapertussis* strain 12822, FR6242 and some other recently collected isolates [24].

To investigate the genomic microevolution of Iranian *B. parapertussis*, reads were mapped against *B. parapertussis* strains 12822 as a reference and a total of 82 SNPs found of which 68 were in coding regions <u>(Supplementary File-2).</u>

From a total of 14 mutations found in intergenic region, ten were in the promoter region of genes including a promoter region of *petA* encoding ubiquinol-cytochrome C reductase iron-sulfur subunit which is a respiratory chain protein that generates an electrochemical potential coupled to ATP synthesis. Another important intergenic mutation was in the promoter region of *bfrE*, the virulence associated gene in *B. pertussis*, encoding probable TonB-dependent receptor for iron transport [3].

From a total of 13 indels (four genic and nine intergenic), four located in coding regions leading to the frameshift mutations of which three were in pseudogenes which may convert them to the active genes (Supplementary File 2). There was no gene insertion or deletion in the genome of the isolate compared to the reference genome.

### Global relationships of *B. parapertussis* isolates

The genetic relationships of the Iranian *B. parapertussis* isolate, IRBP134, with global available isolates was compared (Figure-1) with 102 available human *B. parapertussis* genomes representing seven countries from 1974 to 2018 from multiple studies mainly collected from the United States (Supplementary File-3) [15, 24]. The maximum parsimony tree was constructed using 896 SNPs against *B. parapertussis* strain 12822. From a total of 103 *B. parapertussis* isolates collected during 1974 to 2018, 95 isolates make a distinct lineage from the reference genome by 17 SNPs including two non-synonymous-SNPs in *rnC* and *brkB* encoding ribonuclease III and *Bordetella* serum resistance protein respectively. As discussed previously the nsSNP in *brkB* gene resulted alle variation from 2 in reference genome to 6 (Figure-1). The nsSNP in *rnC* also resulted alle variation from alle 3 to alle 1. The majority of isolates carrying *brkB6* were grouped as clade 1 with 21 (20%) isolates as ancient isolates and clade 2 with 73 isolates (70%) as most recently collected isolates. Most of isolates collected before 2000 were grouped in Clade-1 and share nine common SNPs of which seven located in genes. Furthermore, IRBP134 located in this clade and made a subgroup with two isolates from the USA (FDAARGOS177,1935) and Switzerland (502474-16, 2016) separating from other isolates in Clade 1 with 22 SNPs. FDAARGOS177 which is an FDA standard reference strain and IRBP134 shared 13 SNPs and differed with 2 novel SNPs for IRBP134. The two nsSNPs located in the BPP2476 and BPP3004 encoding hypothetical protein and putative cytochrome C respectively.

## Conclusion

To summarised we identified seven *B. parapertussis* isolates from pertussis cases during 2012-2015. Unlike *B. parapertussis* isolates collected from countries with ACV vaccination that do not express Prn, all Iranian isolates express Prn as one of the major components of ACV vaccine confirming the ACV vaccine pressure on the Prn expression. The global phylogeny analysis showed IRBP134 was grouped with an isolate from the USA and located in the clade 1. To the best of our knowledge, this is the first report investigating the whole genomic features of recently isolated *B. parapertussis*. Our results revealed few mutations showing ongoing genomic adaptation in our isolate. To investigate mutation rate and its effect on the fitness of *B. parapertussis* isolates in Iran, more clinical *B. parapertussis* isolates are needed to be collected from the country to be analysed in terms of genomics or proteomics.

## Acknowledgments

This work was supported financially by Pasteur Institute of Iran grant number 968.

## Conflict of interest

All authors declare that they have no conflict of interest.

## Availability of data and material

The raw reads were submitted to the GeneBank database under the Biosample number SAMN18214790.

## Notes

### Competing Interest Statement

The authors have declared no competing interest.

